# The Global Environment Facility approach for allocating biodiversity funding to countries

**DOI:** 10.1101/2022.12.07.519459

**Authors:** Chris Mcowen, Neil D. Burgess, Neville Ash, Andrea Baquero, Gustavo Fonseca, Mike Harfoot, Craig Hilton-Taylor, Val Kapos, Corinna Ravilious, Catherine Sayor, Oliver Tallowin, Sonja Sabita Teelucksingh, Lauren Weatherdon, Sarah Wyatt

## Abstract

Biodiversity is not evenly distributed across the globe and some areas have greater potential to contribute to biodiversity conservation than others. Whilst there are multiple ways to determine priority areas for conservation, for a global institution like the Global Environment Facility (GEF), the funding mechanism for the Convention on Biological Diversity and the largest multilateral source of funding for developing countries focused on enhancing biodiversity outcomes and promoting sustainable use, it is important to fund the top-ranked countries whilst also ensuring that all eligible countries are able to undertake some biodiversity conservation actions in accordance with the Convention. To this end, the GEF uses the System for Transparent Allocation of Resources (STAR) to allocate funding in separate funding rounds to eligible countries. This country focus means that all prioritization analyses need to be undertaken within that political framework, while also considering the intrinsic patterns in biodiversity that don’t respect national borders. We present the 2018 update of the biodiversity component of GEF-STAR, investigate how the weighting system affects the ranking of countries. We show that top ranked and bottom ranked countries are robust to changes in the weighting of analytical elements, but the weighting can significantly alter the importance of middle ranking countries, affecting their funding allocation. This analysis has been used by the GEF, along with other data, to allocate over $1 billion in biodiversity funding (GEF-7 = $1.2 billion) to improve country and global prospects for conservation. However, this large funding allocation for conservation needs to be set against the vastly larger funding flows that decrease natural values around the world, and the need for systems level change remains evident across the entire planet.

## Introduction

The world is facing a biodiversity crisis potentially leading to an extinction event of geological proportions (Ceballos et al. 2015; Díaz et al. 2019; IPBES 2019). As a response to this crisis, conservation scientists have developed numerous systems to at global to local scales to prioritise conservation efforts, based on different measures of perceived “value”, for example using patterns of biodiversity and threat (Olson & Dinerstein 1998; Mittermeier et al. 2004; Dinerstein et al. 2019; 2020; Jones et al. 2020; Sala et al., in press; Jung et al. in press), or the degree of wilderness, intactness or human footprint (Sanderson et al. 2002; Watson et al. 2018). International NGOs, donor countries, governments, and private philanthropies have, to varying degrees, used such prioritization approaches to inform funding allocation.

However, translating these academic prioritization approaches into the world of donor agencies who work with national governments is challenging and rarely achieved (Beger et al. 2015). There are several reasons for this. First, biologically-defined regions generally cross administrative boundaries, which are subject to overlapping policy and regulatory frameworks making implementation difficult. Second, whilst regional conservation planning clearly provides multiple benefits, decisions, particularly by bilateral and multilateral aid agencies, are made country by country. Therefore, an alternative approach to setting conservation priorities, and seeing them implemented with significant funding resources, is to work at the national level and define biological values in each country with allocated funding for conservation.

The Global Environment Facility (GEF), the financial mechanism of the Convention on Biological Diversity (CBD) and several other multilateral environmental agreements, allocates available funding to eligible countries, including for biodiversity, through an allocation mechanism known as the “System for Transparent Allocation of Resources” (STAR). This system distributes scarce financial resources according to several parameters that include, amongst others, the global environmental benefits that can be generated institutional capacity at a country level, and national income levels, whilst also considering the needs of Least Developed Countries. The systematic and transparent nature of the STAR, including having upper and lower funding limits to reduce large disparities between countries (for non “Least Developed Countries” there is a minimum biodiversity ‘floor’ allocation in GEF-7 of $2 million and “Least Developed Countries” receive a minimum of $3 million), is in keeping with the “leave no one behind” ethos of the Sustainable Development Goals. Further details of this allocation system are found in the STAR Policy (https://www.thegef.org/sites/default/files/council-meeting-documents/EN_GEF.C.54.03.Rev_.01_STAR.pdf)

The original GEF biodiversity global benefits index was also used by other nationally-focused funding agencies, for example it was adopted by USAID’s biodiversity team and used to direct funding flows with small adjustments (USAID, pers comm. Given the national focus of development assistance, country level indices like the GEF-STAR might can also help improve the targeting of development aid between governments, to where it has the greatest potential to conserve biodiversity. This is different funding route to that deployed by NGOs and other agencies that work more on priority regions, sites and species, where the country lens is of secondary consideration.

This paper (1) outlines the method used by the GEF to allocate funds in support of biodiversity conservation at the national level, (2) assesses the method’s sensitivity to changes in the weighting of different input data (3) explores how a nationally focused approach – as used by the GEF – compares with approaches that are based on the importance of biological regions or sites.

## Methods

The methodological approach used to develop the updated GEF-STAR biodiversity index built upon a previous analysis (GEF 2005) which was used by the GEF to allocate resources for GEF-4, GEF-5, and GEF-6 (2006-2018). To maintain a similar funding allocation approach, we retained the various weightings used by the earlier analysis, which are outlined in Annex S1-S3 (consequences of this are discussed later).

For the 2018 analysis we used updated and expanded species data from the IUCN Red List of Threatened Species (www.iucnredlist.org), species and habitat proxy data from the marine realm, and the addition of a sensitivity analysis to assess how country rankings change when scores within the index are adjusted. We also tested the inclusion of freshwater data, but these were eventually dropped as available data were not of sufficient global coverage (see Annex S1).

### Species Patterns

In order to generate biodiversity value layers, we used 30,254 species distribution polygons of all comprehensively assessed species in the IUCN Red List database for birds, mammals, amphibians and a range of marine groups (www.iucnredlist.org). To supplement the relative paucity of marine species we also included nine habitat layers (Annex S1). A raster-based analytical approach (with a 10km grid) was used to complete all subsequent analyses.

### Biodiversity Priority Rankings

The methodology used followed the 2005 approach to ensure comparability of the results with those used in previous GEF funding round, as follows

1. Spatial units were delimited using country/territory codes, using the ISO 3166 classification standard (hereafter ‘countries’). Each country was scored using three attributes for each realm (terrestrial, marine and freshwater): the proportion of the global range of each species present in the country (represented species); the proportion of the global range of each threatened species present in the country (threatened species); the proportion of each distinct terrestrial ecoregion (Dinerstein et al. 2017) present in the country or the proportion of each distinct marine ecoregion (Spalding et al. 2007) present in the country (represented terrestrial and marine ecoregions, respectively) (presented in more detail in Annex S2). The approach used to generate the final country rankings was as follows and was the same as in the 2005 method.

a. Range-size rarity for each threatened species was multiplied by weightings (WT) of 10, 6.7, and 1 for Critically Endangered (CR), Endangered (EN), and Vulnerable (VU).
b. Biodiversity priority ranking was undertaken separately for terrestrial and marine species, as follows: Country Realm Score = WT1 x represented species + WT2 x threatened species + WT3 x + represented ecoregion. Where: WT1=0.65; WT2=0.20; WT3=0.15
c. Marine and terrestrial scores were combined – for each country and used to rank the importance of all countries, as follows: Country Score = 0.75 x Terrestrial Score + 0.25 x Marine Score.
2. To place the relative importance of countries in a global context, the analysis was completed for all countries on Earth (Annex S2), but the results and subsequent funding allocation only relate to the subset of countries that are eligible for GEF support.

A departure from the 2005 approach was to complete a sensitivity analysis of the impacts of the different weighting approaches used (weightings in Annex S4). This entailed varying the weightings of the different components of the score and running 1,000 simulations of each of these options, giving rise to a variety of possible outcomes for the final score per country. Full details of all datasets, methods and decisions made to deliver the country biodiversity rankings are presented in Annex S2, with a detailed workflow in Annex S3.

### Comparison between country biodiversity rankings and biogeographical prioritisation schemes

To facilitate comparison between the country based prioritization approach used here, and the commonly used biogeographically based analyses, we compared GEF STAR results with some of the existing global prioritisations (WWF Global 200 – Olson & Dinerstein 1998 (marine and terrestrial), CI Hotspots –Mittermeier et al. 2004, Key Biodiversity Areas – Eken et al. 2004; Wilderness areas – Sanderson et al. 2002; Jones et al. 2018; Watson et al. 2018)).

## Results

### Biodiversity patterns and priority rankings

Maps of biodiversity value on land and in the marine realm generated for this work (Figures 1a and 1b) illustrate the broad patterns of biodiversity on Earth. These are consistent with other studies, being highest in the tropical regions, especially on wet mountains and along moist coasts, and lowest in colder, flatter and drier areas.

**Figure 1.**
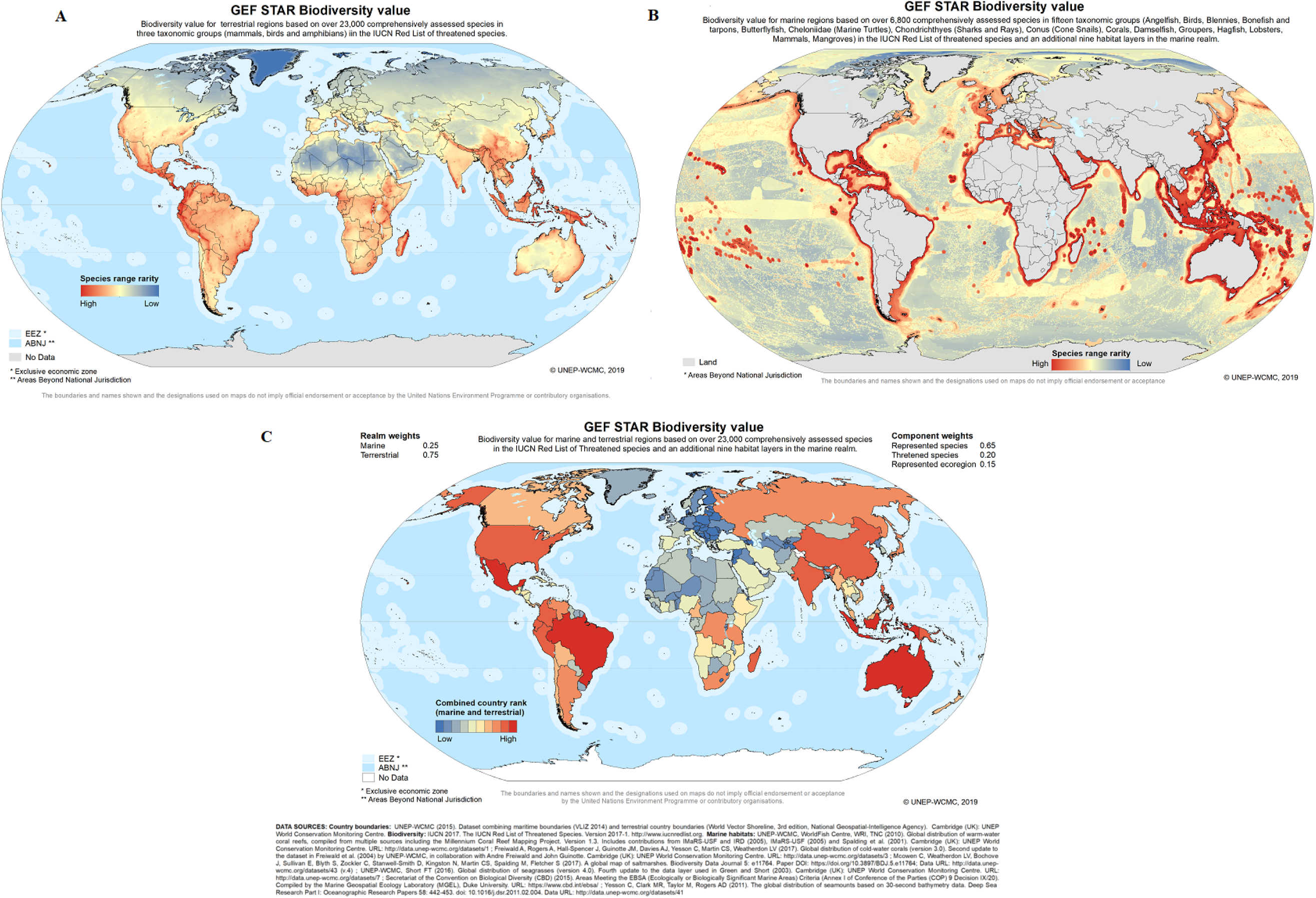
Biodiversity value for a) terrestrial regions; b) marine regions and c) combined using the GEF STAR methodology.

**Figure 2.**
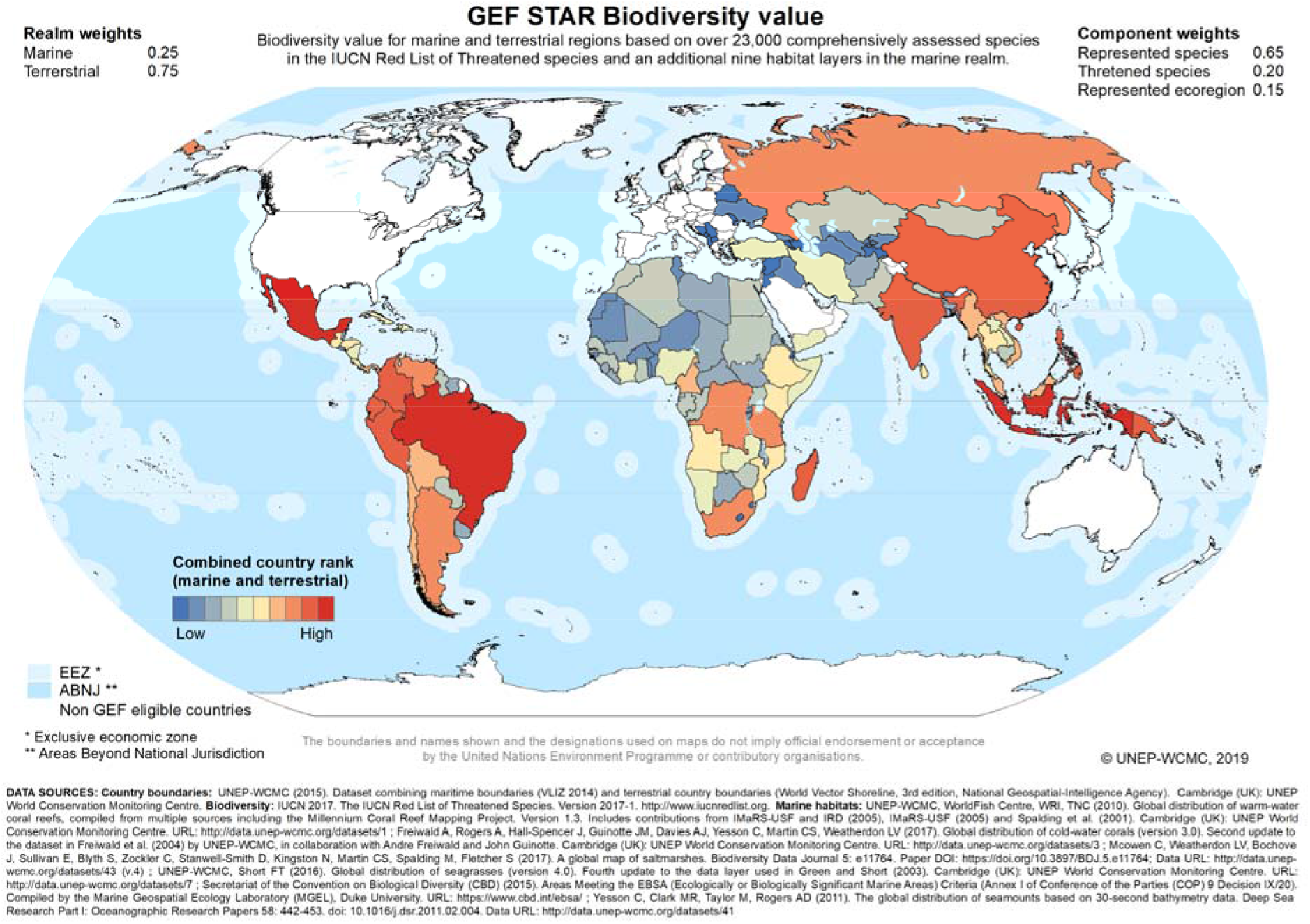
Ranking of countries eligible for GEF funding as calculated using GEF STAR approach

The GEF STAR methodology was then used to combine these terrestrial and marine input data, to produce country-level rankings (Figure 1c). These country rankings for biodiversity then became a key input to the decision-making of the GEF in terms of biodiversity funding allocation. The rankings of all countries on earth are found in Annex S6, along with a note on whether these countries are eligible to receive GEF funding.

### Comparisons between prioritisation schemes

After filtering the global list to contain only those eligible for GEF funding (i.e. the removal of 43% of countries and regions), the GEF STAR approach corresponded best with the Hotspots prioritisation framework. Over half of the countries and regions with KBAs were omitted, 23% of hotspots; 22% of high biodiversity wilderness areas and 17% of global-200 ecoregions (Table 1).

**Table 1.** The top and bottom 20 regions for each prioritisation scheme, filtered to only include those eligible for GEF funding, including the number and percentage of regions on “valued” in the scheme and the degree to which “value” is equally distributed (Gini Index).

|  | Hotspots | Ecoregions | High biodiversity wilderness | Key Biodiversity Areas | GEF STAR |
| --- | --- | --- | --- | --- | --- |
| Top 20 | Brazil | Russian Federation | China | Russia | Indonesia |
|  | Indonesia | Brazil | Russian Federation | Brazil | Brazil |
|  | Mexico | China | Brazil | China | Mexico |
|  | Chile | Indonesia | Argentina | Algeria | Colombia |
|  | Kiribati | DRC | Kiribati | Libya | China |
|  | China | PNG | Indonesia | Niger | Madagascar |
|  | Ethiopia | Tanzania | Venezuela | Mauritania | Peru |
|  | Madagascar | Colombia | Bolivia | Mali | India |
|  | Micronesia | Peru | Peru | Chad | Philippines |
|  | Myanmar | Turkey | Algeria | Indonesia | PNG |
|  | Philippines | Mongolia | South Africa | Egypt | Ecuador |
|  | Cook Islands | Argentina | India | Colombia | Venezuela |
|  | Marshall Islands | Venezuela | Tanzania | Venezuela | South Africa |
|  | Turkey | Myanmar | DRC | Peru | Argentina |
|  | Thailand | Mexico | Kazakhstan | Botswana | Tanzania |
|  | India | Philippines | Turkey | Central African Republic | Russia |
|  | Colombia | Madagascar | Ethiopia | Angola | DRC |
|  | Iran | Mozambique | Botswana | Mongolia | Malaysia |
|  | Viet Nam | Angola | Chad | Sudan | Bolivia |
|  | Malaysia | Thailand | Myanmar | Suriname | Chile |
| Bottom 20 | Burundi | Equatorial Guinea | Gambia | Jamaica | Syria |
|  | Barbados | Rwanda | Bosnia and Herzegovina | Palau | Azerbaijan |
|  | Rwanda | Ethiopia | Djibouti | Haiti | Tajikistan |
|  | Grenada | Timor-Leste | Burundi | Marshall Islands | Djibouti |
|  | Dominica | Côte d'Ivoire | Samoa | Dominica | Albania |
|  | Bangladesh | Bosnia and Herzegovina | Trinidad and Tobago | Kiribati | Niue |
|  | Zimbabwe | Macedonia | Guyana | Timor-Leste | Armenia |
|  | Montenegro | Nigeria | Seychelles | Dominican Republic | Lesotho |
|  | Togo | Gambia | Mauritius | Antigua and Barbuda | Nauru |
|  | Saint Lucia | Lebanon | Republic of Moldova | Togo | Bosnia and Herzegovina |
|  | Macedonia | Serbia | Sao Tome and Principe | Mauritius | Belarus |
|  | Bosnia and Herzegovina | Moldova | Saint Vincent and the Grenadines | Barbados | Swaziland |
|  | South Sudan | Montenegro | Saint Lucia | Iraq | Serbia |
|  | Saint Kitts and Nevis | Vanuatu | Antigua and Barbuda | Saint Kitts and Nevis | Gambia |
|  | Equatorial Guinea | Jordan | Dominica | Albania | Saint Kitts and Nevis |
|  | Benin | South Sudan | Saint Kitts and Nevis | Bhutan | Jordan |
|  | Sao Tome and Principe | Iraq | Niue | Eritrea | Montenegro |
|  | Trinidad and Tobago | Sao Tome and Principe | Grenada | Equatorial Guinea | Lebanon |
|  | Serbia | Mali | Maldives | Grenada | Macedonia |
|  | Uruguay | Tonga | Barbados | Saint Lucia | Moldova |
| Total number | 121 | 90 | 103 | 100 | 141 |
| % of total | 86 | 64 | 73 | 71 | 100 |
| GINI index | 0.73 | 0.84 | 0.73 | 0.93 | 0.69 |

Fifty countries (40%) are highly valued by all prioritisation schemes. These tend to be large, diverse countries, such as Indonesia and Brazil). Further to this, 62 countries (44%) are valued by four schemes; 17 countries (12%) by three, and five countries (4%) by two. Of the five countries valued by only two schemes (Burkina Faso, Tuvalu, Lesotho, Belarus and Nauru), three are valued only by the GEF STAR and KBA frameworks and two are valued only by the hotspots and GEF STAR approach. It is apparent that the top-ranking countries eligible for GEF funding have disproportionately high “value” for both the both the terrestrial and marine realm scores using the GEF STAR approach.

### Sensitivity of the GEF STAR approach to different weightings

A sensitivity analysis of the influence of weighting decisions inherent in the GEF methodology (see Annex S2 and S3), shows that countries at the top and - to a lesser extent – at the bottom of the list are relatively stable in terms of their ranked position. In comparison, the ranking of countries in the middle of the table alters considerably depending on the weightings used, as illustrated by the top 20, middle 20 and bottom 20 countries (Figure 3).

**Figure 3.**
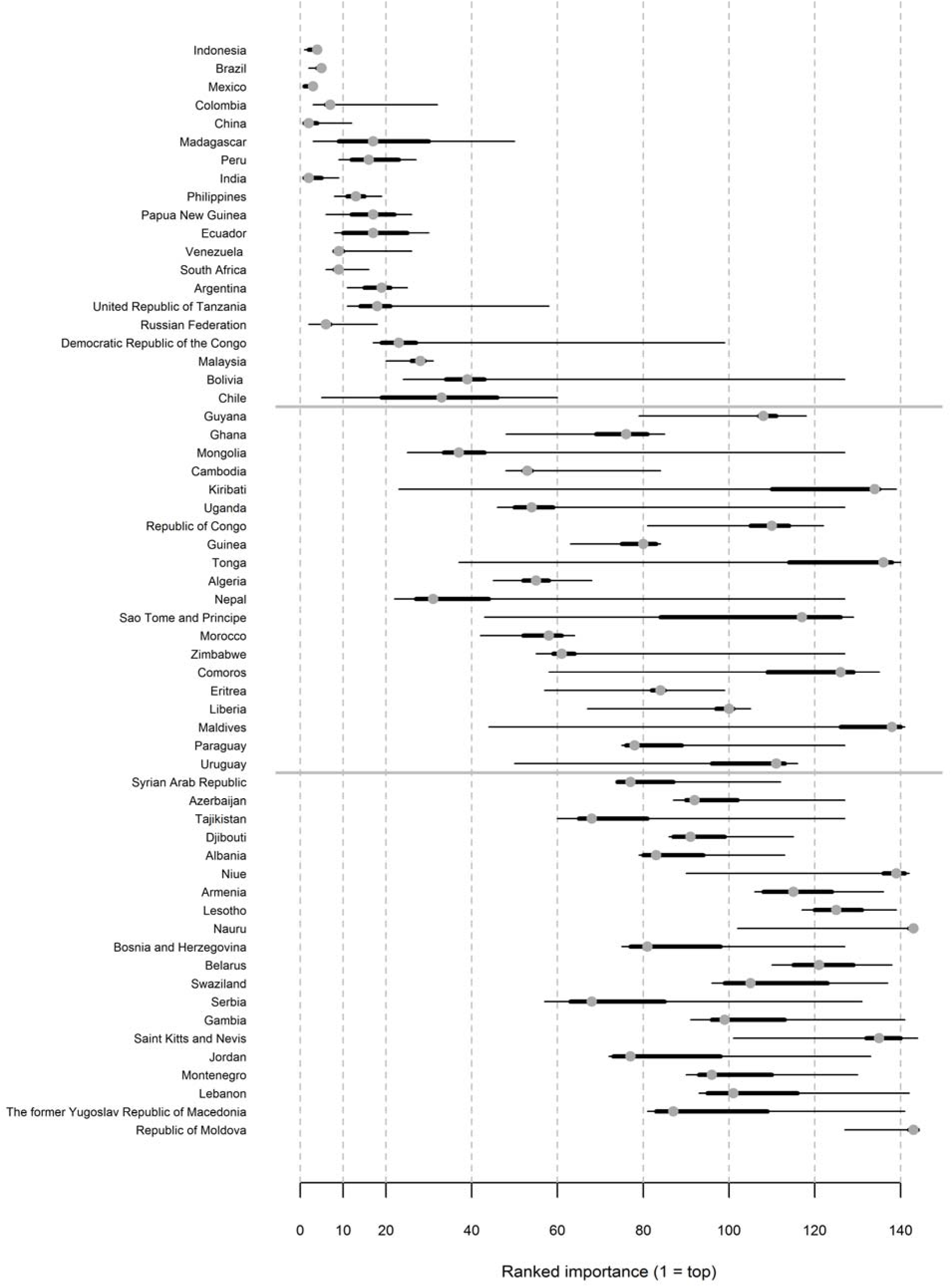
Sensitivity analysis for top and bottom ranked and selected middle ranking countries as affected by changing the weightings of different parameters in the GEF STAR Index

## Discussion

This paper illustrates, using a real-world example from the GEF, the challenges faced by donor agencies when determining how to allocate scarce financial resources to countries which are the unit of operation.

### Benefits and challenges of the GEF funding allocation model

A number of GEF-eligible countries are of particular importance in our analysis. For example, Indonesia is at the top of the rankings for both terrestrial and marine biodiversity, whereas others such as Brazil are critical mainly for their terrestrial importance. Funds from the GEF are allocated based on these scores, hence countries with large amounts of biodiversity will receive relatively larger allocations, which makes sense as an approach to ensure maximum conservation gain for the available funding.

However, preventing biodiversity loss and extinction requires action everywhere, irrespective of a country’s as there is unique biodiversity in every country; therefore,. allocating all or the vast majority of GEF funding to a handful of ‘megadiverse or hotspot’ countries would not be achieve the goal of halting biodiversity loss. The GEF funding allocation framework takes account of this challenge as the scores are used to allocate funds. Countries in the middle of the rankings receive more funding than lower ranked countries. However, it is recognized that even countries with low scores have biodiversity importance as even with less biodiversity there is likely different and unique biodiversity not found in the high diversity countries, and these countries therefore also receive funding, albeit in smaller amounts. As such an important benefit of the GEF ‘STAR’ approach is that *every* eligible country receives financing and can deploy these funds to make meaningful impacts. This is especially true in developing countries, which are typically underfunded, and where modest financial investment can result in significant changes. Allocated funding is also often used to strengthen the capacity of, and communication between, national environmental organisations, leading to more widespread, sustainable and longer-term impact. Furthermore, funding can be used to develop national environmental strategies that cut across departments and sectors and can bring about the transformative changes required to address systemic drivers of biodiversity loss, for example through mainstreaming or restoration agendas.

### Comparison of GEF financial support with other funding flows

While it is important to ensure that biodiversity conservation funding is directed to places of high biodiversity importance, and the GEF is the largest conservation funding source available, it is also important to remember that conservation funding is very limited in comparison to larger economic forces. The GEF-7 biodiversity funding for four years is $1.2 billion; however, in 2016 Americans spent approximately $66 billion on pet food and $20 billion on ice cream. The GEF funding therefore remains a tiny proportion of global financial flows and much of the funding being deployed around the world drives the loss of nature. As such, GEF funding cannot be expected to solve the biodiversity crisis alone and needs to leverage other resources. Recent research (Waldron et al. 2013; 2020) has also suggested that there is a huge economic return from conservation in investment and implementation, so the funds from the GEF are likely to be leveraging further economic benefits.

Finally, as the world seeks to negotiate and agree a new agreement for the next 10 years under the Convention on Biological Diversity, the GEF will clearly remain a key player in the delivery of aspects of that agreement. The work presented here shows how funding flows can be channelled and other work (for example Leclère et al. 2020, SCBD 2020) has shown some of the key things that need to be done to reverse natures decline. Taken as a part of a package of interventions the work of the GEF can lead the world towards more sustainable “nature positive future”.

## Supporting information

Annex S1

Annex S2

Annex S3

## Acknowledgements

We would like to thank all the data providers to the IUCN Red List as without their efforts analyses such as these would be impossible. Marine habitat data comes from Ocean+ system managed at UNEP-WCMC. Ecoregions 2017 come from Resolve in the USA. Other data we used here have been made freely available by various agencies, we thank them and their funders. We also thank the GEF Secretariat for funding.

## Notes

### Competing Interest Statement

The authors have declared no competing interest.

