## Supplementary material for "The Global Environment Facility approach for allocating biodiversity funding to countries": Annex S1

### **Annex S1: Datasets**

All the datasets used in this analysis are global in extent, and their sources are provided in this Annex.

#### **Country boundaries**

| **Dataset** | **Version / Date** | **Organisation** | **Update frequency** | **Citation and hyperlink** |
| --- | --- | --- | --- | --- |
| Maritime and Terrestrial Country Boundaries^[1]^ | 2015 | UN Environment World Conservation Monitoring Centre | Data are updated in intervals that are uneven in duration | UNEP-WCMC (2015). Dataset combining maritime boundaries (VLIZ 2014) and terrestrial country boundaries (World Vector Shoreline, 3rd edition, National Geospatial-Intelligence Agency). Cambridge (UK): UNEP World Conservation Monitoring Centre. |

**Justification**

- **Maritime and Terrestrial Country Boundaries:** This is the only freely available data set that combines maritime and terrestrial boundaries in one layer

#### **Ecoregions**

| **Dataset** | **Version / Date** | **Organisation** | **Update frequency** | **Citation and hyperlink** |
| --- | --- | --- | --- | --- |
| Marine Ecoregions and Pelagic Provinces of the World | 2012 | The Nature Conservancy (TNC) | Data are updated in intervals that are uneven in duration | The Nature Conservancy (2012). Marine Ecoregions and Pelagic Provinces of the World. GIS layers developed by The Nature Conservancy with multiple partners, combined from Spalding et al. (2007) and Spalding et al. (2012). Cambridge (UK): The Nature Conservancy. DOIs: 10.1641/B570707 <http://data.unep-wcmc.org/datasets/38> |
| Terrestrial ecoregions of the world | 2017 | Resolve (USA) – updating the previous dataset of the World Wildlife Fund (WWF) | Data are updated in intervals that are uneven in duration | E. Dinerstein et al., Bioscience 10.1093/biosci/bix014 (2017) <https://academic.oup.com/bioscience/article-lookup/doi/10.1093/biosci/bix014> |
| Freshwater Ecoregions of the World^[2]^ | 2008 | World Wildlife Fund (WWF) and The Nature Conservancy TNC) | Data are updated in intervals that are uneven in duration | Abell et al. (2008). Freshwater Ecoregions of the World: A New Map of Biogeographic Units for Freshwater Biodiversity Conservation. BioScience 58(5):403-414.  <https://doi.org/10.1641/B580507> , <https://academic.oup.com/bioscience/article-lookup/doi/10.1641/B580507> |

**Justification**

- **Marine Ecoregions and Pelagic Provinces of the World:** This dataset combines two separately published datasets: the “Marine Ecoregions Of the World” (MEOW) and the “Pelagic Provinces Of the World” (PPOW). The MEOW dataset shows a biogeographic classification of the world's coastal and continental shelf waters, following a nested hierarchy of realms, provinces and ecoregions. The regions aim to capture generic patterns of biodiversity across habitats and taxa, with regions extending from the coast (intertidal zone) to the 200 m depth contour (extended beyond these waters out by a 5km buffer). The PPOW dataset shows a biogeographic classification of the surface pelagic (i.e. epipelagic) waters of the world's oceans. It describes 37 pelagic provinces of the world, nested into four broad realms. A system of seven biomes are also identified ecologically, and these are spatially disjoint but united by common abiotic conditions, thereby creating physiognomically similar communities. This is the only global dataset that is consistent across the many marine realms and coastal zones.
- **Terrestrial ecoregions of the world:** This map offers the most comprehensive coverage, and follows a classification framework that builds on existing biogeographic knowledge, at a detailed level of biogeographic resolution. The dataset used is an update of the original database developed by WWF in 2001 and corrects several errors in the data and updates some regions to take on board new knowledge.
- **Freshwater Ecoregions of the World^[3]^:** This is the only global dataset available for freshwater ecoregions

#### **Marine habitats and areas of biodiversity importance**

| **Dataset** | **Version / Date** | **Update frequency** | **Organisation** | **Citation and hyperlink** |
| --- | --- | --- | --- | --- |
| Global Distribution of Mangroves USGS | 2011 | Corrections are made on an ad-hoc basis | UN Environment World Conservation Monitoring Centre (UNEP-WCMC) | Giri C, Ochieng E, Tieszen LL, Zhu Z, Singh A, Loveland T, Masek J, Duke N (2011). Status and distribution of mangrove forests of the world using earth observation satellite data (version 1.3, updated by UNEP-WCMC). Global Ecology and Biogeography 20: 154-159. doi: [10.1111/j.1466-8238.2010.00584.x](http://dx.doi.org/10.1111/j.1466-8238.2010.00584.x) . Data URL: <http://data.unep-wcmc.org/datasets/4> |
| Global Distribution of Coral Reefs | 2010 | Corrections are made on an ad-hoc basis | UN Environment World Conservation Monitoring Centre (UNEP-WCMC) | UNEP-WCMC, WorldFish Centre, WRI, TNC (2010). Global distribution of warm-water coral reefs, compiled from multiple sources including the Millennium Coral Reef Mapping Project. Version 1.3. Includes contributions from IMaRS-USF and IRD (2005), IMaRS-USF (2005) and Spalding et al. (2001). Cambridge (UK): UNEP World Conservation Monitoring Centre. URL: <http://data.unep-wcmc.org/datasets/1> |
| Global Distribution of Cold-water Corals | 2017 | Data are updated in intervals that are uneven in duration | UN Environment World Conservation Monitoring Centre (UNEP-WCMC) | Freiwald A, Rogers A, Hall-Spencer J, Guinotte JM, Davies AJ, Yesson C, Martin CS, Weatherdon LV (2017). Global distribution of cold-water corals (version 3.0). Second update to the dataset in Freiwald et al. (2004) by UNEP-WCMC, in collaboration with Andre Freiwald and John Guinotte. Cambridge (UK): UNEP World Conservation Monitoring Centre. URL: <http://data.unep-wcmc.org/datasets/3> |
| Global Distribution of Saltmarshes | V. 4  2017 | Data are updated in intervals that are uneven in duration. | UN Environment World Conservation Monitoring Centre (UNEP-WCMC) | Mcowen C, Weatherdon LV, Bochove J, Sullivan E, Blyth S, Zockler C, Stanwell-Smith D, Kingston N, Martin CS, Spalding M, Fletcher S (2017). A global map of saltmarshes. Biodiversity Data Journal 5: e11764. Paper DOI: <https://doi.org/10.3897/BDJ.5.e11764>; Data URL: <http://data.unep-wcmc.org/datasets/43> (v.4) |
| Global Distribution of Seagrasses | V. 4  2016 | Data are updated in intervals that are uneven in duration. | UN Environment World Conservation Monitoring Centre (UNEP-WCMC) | UNEP-WCMC, Short FT (2016). Global distribution of seagrasses (version 4.0). Fourth update to the data layer used in Green and Short (2003). Cambridge (UK): UNEP World Conservation Monitoring Centre. URL: <http://data.unep-wcmc.org/datasets/7> |
| Global Distribution of Ecologically or Biologically Significant Marine Areas | 2015 | Data are updated in intervals that are uneven in duration. | Convention on Biological Diversity (CBD) Secretariat | Secretariat of the Convention on Biological Diversity (CBD) (2015). Areas Meeting the EBSA (Ecologically or Biologically Significant Marine Areas) Criteria (Annex I of Conference of the Parties (COP) 9 Decision IX/20). Compiled by the Marine Geospatial Ecology Laboratory (MGEL), Duke University. URL: https://www.cbd.int/ebsa/ |
| Global Distribution of Cold Seeps | 2010 | Data are not being updated. | National Oceanography Centre | Baker MC, Ramirez-Llodra E, Perry D (2010). ChEssBase: an online information system on species distribution from deep-sea chemosynthetic ecosystems. Version 3. Chemosynthetic Ecosystem Science (ChEss) project. Southampton (UK): National Oceanography Centre. URL: [www.noc.soton.ac.uk/chess](http://www.noc.soton.ac.uk/chess) |
| Global Distribution of Hydrothermal Vent Fields | v. 3.3  2015 | Data are updated in intervals that are uneven in duration. | Woods Hole Oceanographic Institution | Beaulieu, S., Joyce, K., Cook, J. and Soule, S.A (2015). InterRidge Vents Database Version 3.3. Woods Hole Oceanographic Institution (2015). <http://vents-data.interridge.org> |
| Global Distribution of Seamounts and Knolls | 2011 | Data are not being updated | Zoological Society of London (ZSL) | Yesson C, Clark MR, Taylor M, Rogers AD (2011). The global distribution of seamounts based on 30-second bathymetry data. Deep Sea Research Part I: Oceanographic Research Papers 58: 442-453. doi: [10.1016/j.dsr.2011.02.004](http://dx.doi.org/10.1016/j.dsr.2011.02.004). Data URL: <http://data.unep-wcmc.org/datasets/41> |

**Justification**

- **Global Distribution of Mangroves USGS:** Mangroves serve as valuable nurseries and habitats for many commercially important and threatened and endangered species. This is the most recent, globally consistent map of mangrove distribution.
- **Global Distribution of Coral Reefs:** Coral reefs support more species per unit area than any other marine environment, including about 4,000 species of fish, 800 species of hard corals and hundreds of other species. This is the most recent, globally consistent map of coral distribution.
- **Global Distribution of Cold-water Corals:** Cold water corals act as important habitats for many species, including squat lobsters, starfish, sea urchins, anemones and sponges and are important breeding grounds for many commercially important fish. Although the number of fish species is relatively low, the number of invertebrate species on reefs in the Northeast Atlantic Ocean have been found to be as high as that found in shallow-water tropical reefs. This is the most recent, globally consistent map of cold-water coral distribution.
- **Global Distribution of Saltmarshes:** Saltmarshes are highly productive, often overlooked ecosystems that are rich in biodiversity, consisting of a variety of plant species that provide important habitats for a wealth of fauna. This is the first global map of salt marsh distribution.
- **Global Distribution of Seagrasses:** Seagrasses are a vital part of the marine ecosystem and provide food, habitat, and nursery areas for numerous vertebrate and invertebrate species. The vast biodiversity and sensitivity to changes makes seagrasses an important species to help determine the overall health of coastal ecosystems. This is the most recent, globally consistent map of seagrass distribution.
- **Global Distribution of Ecologically or Biologically Significant (marine) Areas:** these areas are the result of a description process convened by the Secretariat of the Convention on Biological Diversity (CBD). There are seven criteria for identifying EBSAs, of which one or more must be applicable: Uniqueness or rarity; Special importance for life history stages of species; Importance for threatened; endangered or declining species and/or habitats; Vulnerability; fragility; sensitivity or slow recovery; Biological productivity; Biological diversity; and Naturalness.
- **Global Distribution of Cold Seeps:** Cold seeps are shallow areas on the ocean floor where gases percolate through underlying rock and sediment layers and emerge on the ocean bottom. These areas support a wide diversity of life and due to their unique environment they support a number of endemic species. This is the most recent, globally consistent map of cold seep distribution.
- **Global Distribution of Hydrothermal Vent Fields:** Deep-sea hydrothermal vents form as a result of volcanic activity on the ocean floor and support thriving communities of shrimp, crabs, giant tubeworms, clams, slugs, anemones, and fish, with a biomass with a biomass equivalent to that of a rainforest. This is the most recent, globally consistent map of hydrothermal vent distribution.
- **Global Distribution of Seamounts and Knolls:** Seamounts and knolls are underwater mountains. Often referred to as ocean oases, they provide important aquatic habitat for benthic invertebrates, deep-sea fish, and marine predators. Hotspots for biodiversity, seamounts support many commercially important marine species. This is the most recent, globally consistent map of seamounts and knolls distribution.

#### **Threat and pressure**

| **Dataset** | **Realm** | **Version / Date** | **Update frequency** | **Citation and hyperlink** |
| --- | --- | --- | --- | --- |
| Spatial and temporal changes in cumulative human impacts on the world’s ocean | Marine | 2015 | Data are updated in intervals that are uneven in duration. | Halpern, B. S. *et al.* Spatial and temporal changes in cumulative human impacts on the world’s ocean. Nat. Commun*.* 6:7615 doi: [10.1038/ncomms8615](https://www.nature.com/articles/ncomms8615) (2015). |
| Global terrestrial Human Footprint maps for 1993 and 2009 | Terrestrial | 2016 | Data are updated in intervals that are uneven in duration. | Venter, O., Sanderson, E.W., Magrach, A., Allan, J.R., Beher, J., Jones, K.R., Possingham, H.P., Laurance, W.F., Wood, P., Fekete, B.M., Levy, M.A., Watson, J.E.M., 2016a. Global Human Footprint maps for 1993 and 2009. Sci. Data 3, 10067 |
| Global threats to human water security and river biodiversity^[4]^ | Freshwater | 2010 | Data are updated in intervals that are uneven in duration. | Vörösmarty, C. J., McIntyre, P. B., Gessner, M. O., Dudgeon, D., Prusevich, A., Green, P., & Davies, P. M. (2010). Global threats to human water security and river biodiversity. Nature, 467(7315), 555-561. DOI: 10.1038/nature09440 |

**Justification**

- **Spatial and temporal changes in cumulative human impacts on the world’s ocean:** Quantifying and mapping local- and global-scale stressors in a standardized, comparable manner offers a powerful means to assess the spatial pattern of individual human pressures, as well as their total impact on natural systems across highly variable geographies. This dataset maps the cumulative impact of 19 different types of anthropogenic stress on 20 global marine ecosystem types using best available global-scale data as of 2013.
- **Global terrestrial Human Footprint maps for 1993 and 2009:** Human pressures on the environment are the actions taken by humans with the potential to harm nature. Cumulative pressure mapping measures the breadth of these pressures by coupling top-down remote sensing of land cover change with data on additional human pressures collected ‘bottom-up’ through systematic surveys and modelling. This map represents the most current information of its type.
- **Global threats to human water security and river biodiversity**^[5]^**:** Many stressors threaten human water security and biodiversity through similar pathways, as for pollution, but they also influence water systems in distinct ways. This is the first globally consistent map of cumulative pressures upon freshwater biodiversity.

#### **Species**

Species ranges come from a database maintained by IUCN^[6]^.

| **IUCN Terrestrial Species Groups** | **IUCN Marine species groups** | **Freshwater** species groups^[7]^ |
| --- | --- | --- |
| Amphibians | Angelfish | Crabs |
| Birds | Birds | Crayfish |
| Chamaeleonida (Chameleons) | Blennies | Shrimps |
| Crocodylia (Crocodiles/Alligators) | Bonefish and tarpons | Sturgeons |
| Mammals | Butterflyfish |  |
|  | Cheloniidae (Marine Turtles) |  |
|  | Chondrichthyes (Sharks and Rays) |  |
|  | Conus (Cone Snails) |  |
|  | Corals |  |
|  | Damselfish |  |
|  | Groupers |  |
|  | Hagfish |  |
|  | Lobsters |  |
|  | Mammals |  |
|  | Mangroves |  |
|  | Seabreams and porgies |  |
|  | Pufferfish |  |
|  | Sea cucumbers |  |
|  | Seagrasses |  |
|  | Sea snakes |  |
|  | Sturgeonfish, tangs and unicornfish |  |
|  | Tunas and billfishes |  |
|  | Wrasse |  |

1. This dataset is one that has been developed by UN Environment World Conservation Monitoring Centre (UNEP-WCMC) by combining two existing datasets. It provides political boundaries for the terrestrial and marine realms, including territorial waters and Exclusive Economic Zones (EEZ) for the latter.
2. Used in exploratory analysis only
3. Used in exploratory analysis only
4. Used in exploratory analysis only
5. Used in exploratory analysis only
6. IUCN 2017. The IUCN Red List of Threatened Species. Version 2017-1. <<http://www.iucnredlist.org>>.
7. Used in exploratory analysis only
