## Supplementary material for "The Global Environment Facility approach for allocating biodiversity funding to countries": Annex S2

**Methodological approach**

The final method characterises each country using three main scores— represented species, threatened species and represented ecoregions—each of which is described in detail below. These were calculated in a consistent manner across terrestrial and marine realms, using the following steps that are described more fully in the following sections, and with a detailed technical workflow found in Annex 2: Technical Workflow.

**Step 1: Prepare represented species and threatened species layers (separately for each realm)**

Preparing and processing the species datasets for steps 3-5 below is a time-consuming task that requires considerable computer processing resources. Several python scripts and processing models in ArcGIS have been generated to increase the efficiency of this task in this analysis and for any future revisions/updates. The data covered species from all taxonomic groups that have been comprehensively assessed (see Section 3.5 ‘Species’ in Annex for details of the taxonomic groups included), resulting in a database of 23,442 species in the terrestrial realm and 6,812 in the marine (freshwater species were excluded for reasons explained above).

**Step 2: Prepare marine habitat datasets (marine realm only)**

Additionally, for the marine realm, the distribution of important habitats and biologically important areas were considered in the represented species score (see Section 3.3 ‘Marine habitats and areas of biodiversity importance’ in Annex: Datasets for details of habitats). Each marine habitat layer was converted to a raster, based on the proportion of a cell occupied by the habitat, ensuring that the amount of habitat in each 10km grid cell is recorded.

**Step 3: Create the scores for represented species (separately for each realm)**

Each 10km grid cell was scored for range-size rarity for each species (the proportion of the species’ global range the cell represents; i.e. 1/range size) and given a total score by summing scores across all the species potentially occurring in it. Each represented species contributes to the component based on the proportion of its global range within each 10km grid cell.

**Step 4: Create the refined marine scores for represented species (marine realm only)**

For each habitat, the proportion of area relative to the global extent is calculated for each grid cell. This approach is consistent with how individual species were scored (as above). The marine habitats datasets function as proxies for species occurring within these habitats. In most cases, such species were not included in the IUCN Red List species data, particularly those of habitats such as seamounts, seeps, vents and cold-water corals. For consistency in the marine approach, coastal habitats, e.g. mangroves, were also included. Each marine habitat was in effect treated as an additional species and combined with the marine represented species score generated from the IUCN Red List species data.

**Step 5: Create the scores for threatened species (separately for each realm)**

This score considered the subset of species from the represented species score that are assessed as threatened^[1]^—i.e. Critically Endangered (CR), Endangered (EN), or Vulnerable (VU)—on the IUCN Red List. The range-size rarity for each threatened species was multiplied by weightings of 10, 6.7, and 1 for CR, EN, and VU, respectively. These weightings are the same as those used in the original methodology. These weighted range-size rarity values were then summed in each grid cell. Each threatened species therefore contributed to the component score based on the proportion of its global range within each 10km grid cell, weighted based on its relative extinction risk.

**Step 6: Prepare the country ecoregion datasets (CECs) and create the scores for representative ecoregions (separately for each realm);**

This analytical step involved the preparation of a CEC layer, by overlaying biologically determined ecoregion maps with politically-determined country boundaries (see Sections 1 and 2 in Annex: Datasets for a list of the datasets used). Each realm had a distinct set of CECs based on the realm-specific ecoregions layer. The 828 ecoregions making up the terrestrial realm for GEF-eligible countries encompass 1,675 CECs. For the marine realm, 260 ecoregions encompass 8,282 CECs.

For each ecoregion, an equivalent measure to the range-size rarity score for species was calculated. In other words, each ecoregion was represented proportionally to its global extent, i.e. 1/global extent per 10km grid cell. This meant that when summed at CEC level, each ecoregion contributed to the **represented ecoregion score** based on the proportion of its global extent within each CEC. High scores for ‘represented ecoregions’ occur in CECs containing large proportions of the ecoregion’s area. When summed at a country level, the score reflects both the number of ecoregions in the country and the number of high and low scores for its CECs.

**Step 7: Generate country level scores for each of the 3 component scores (separately for each realm) and export to excel.**

This analytical step summed the pixel level scores for each country, separately for each of the components for each of the realms. Although interim tables were generated at CEC and country level, the final tables were only required by the GEF at country level only, noting however that even at the country level CEC’s are still considered through the representative ecoregion component score. Due to the approach taken, i.e. performing the calculations at 10km pixels, it is possible to aggregate pixels at various spatial units (e.g. CEC, Country, and EEZ).

**Step 8: Normalise and weight the 3 component scores for each country and combine into a composite score for each country (separately for each realm)**

The penultimate analytical step scaled the composite score for each country. Each component score was normalized from 0-100 and then multiplied by the defined component weight as shown in the following equation:

*Country Biodiversity Realm Score = WT1 x Represented Species + WT2 x Threatened Species + WT3 x + Represented Ecoregion*

*Where*

*WT1=0.65; WT2=0.20; WT3=0.15^[2]^*

Aligning with the previous approach, larger weights are given to species attributes as these are characterised with greater certainty. Furthermore, threatened species are given additional weight through the inclusion of the ‘threatened species’ component score, which accounts for the relative extinction risk of species.

**Step 9: Combine the realm scores (in excel) to get the final GBI_BD_ for each country**

The last analytical step (Step 9) combined the realm scores. The Country Biodiversity Realm Score generated in Step 8 were summed together applying additional realm weights.

*GBI_BD_ = WT x Terrestrial Score + WM x Marine Score*

*Where*

*WT=0.75; WM=0.25^3^*

### Caveats

- Although all IUCN Red List species range data for comprehensively assessed taxonomic groups have been included in the analysis, these represent only part of all biodiversity known to occur globally, and are biased towards vertebrates. This means that our analysis may have reduced the apparent importance of some countries that have ecoregions with rich floral and/or invertebrate diversity, but relatively poor vertebrate fauna.
- It is also important to note that the IUCN Red List spatial data captures the total range of a given species (‘extent of occurrence’), rather than precise areas where that species is confirmed to be present. This is particularly relevant for species with broad ranges and limited documentation of detailed occurrences within specific habitats.
- The species and habitat datasets used for this analysis are known to have limited data coverage in data-poor regions, and are therefore biased towards data-rich regions. Thus, limited data availability may inadvertently lead to downgrading of countries, particularly developing countries that are biodiverse but have limited capacity and/or resources to document this biodiversity through comprehensive monitoring initiatives.
- For those global terrestrial and marine spatial datasets that do exist, it can be difficult to ensure consistency across regions. Many global datasets are collated from multiple sources, with contributing datasets often differing in spatial resolution, quality, and methodological approach. Furthermore, countries with greater financial or technological capacities are likely to have more detailed maps and inventories of the species present. Consequently, these datasets should be interpreted as high-level indications of the approximate locations of biodiversity, rather than systematic, homogenous and complete records of biodiversity.

1. Unlike the previous analysis, Extinct in the Wild species were excluded from this analysis. This resulted in 9 species being removed from the analysis.
2. NB. A sensitivity analysis was conducted testing various weights for each component and realm. The results were found to be robust to changes and the final weights used were defined as per recommendations from the GEF.
