## Supplementary figures and images for "The Global Environment Facility approach for allocating biodiversity funding to countries"

### Annex S3

#### **Annex S3: Technical Workflow**

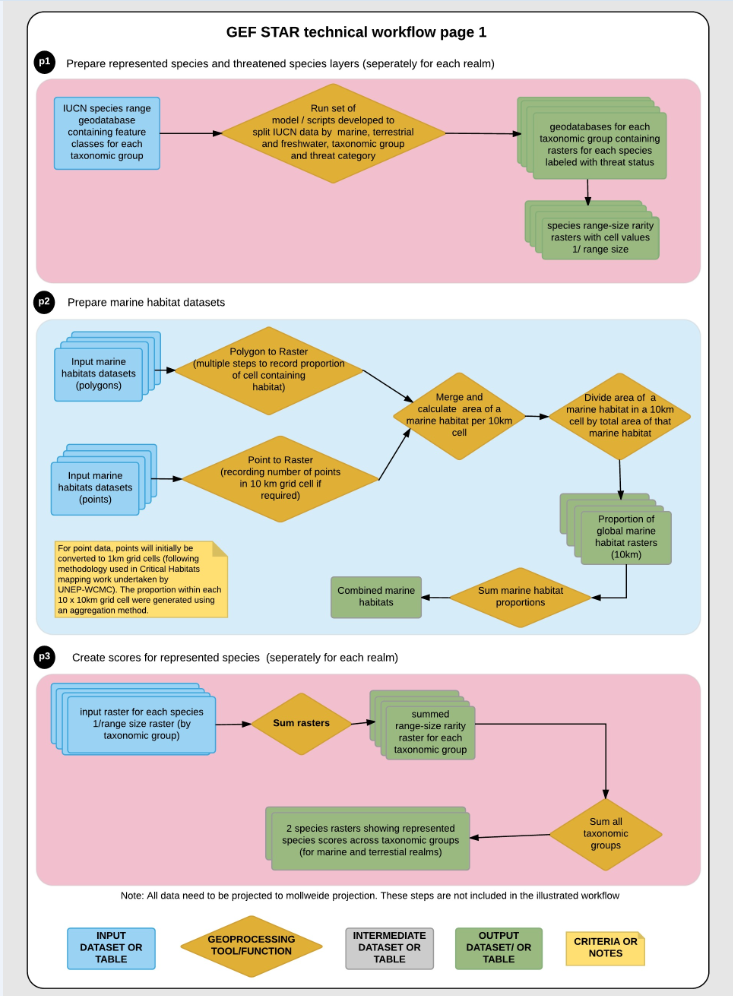

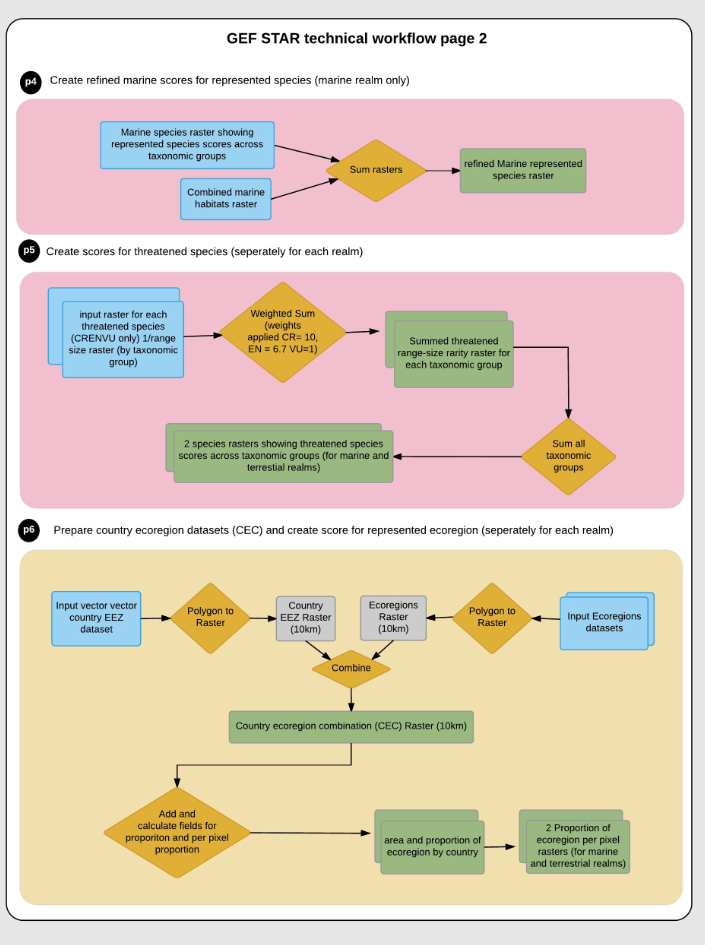

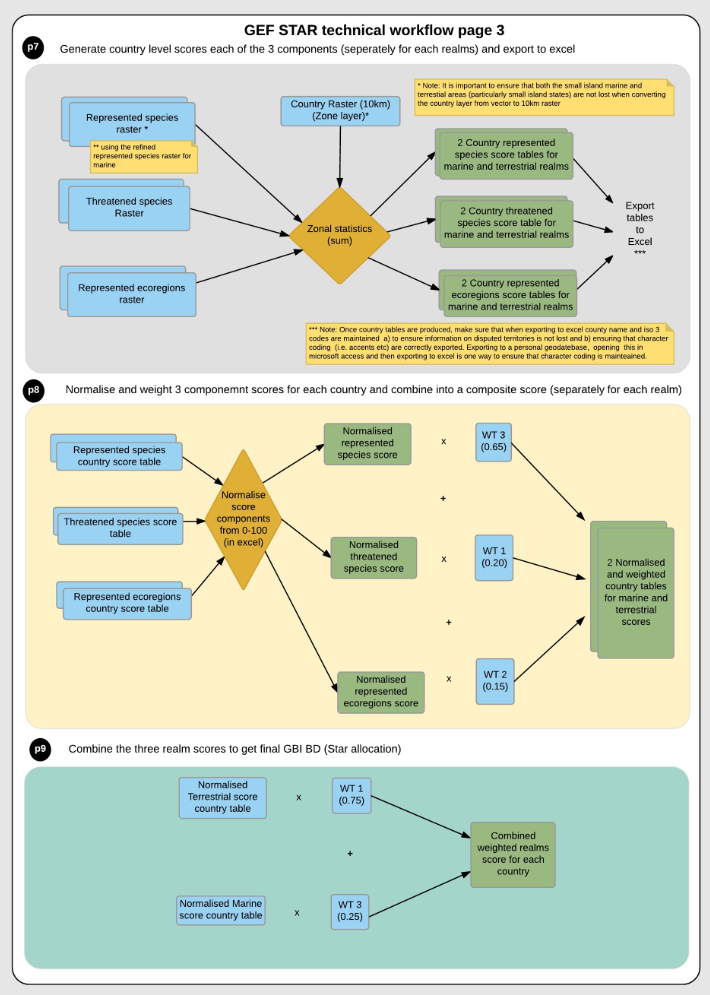
